# Dynamic optimization of chemo–immunotherapy sequencing reveals phenotype-dependent regimens across tumor immune microenvironments

**DOI:** 10.1101/2025.11.26.690636

**Authors:** Kai Gong, Tong Lu, Xu Wang, Xinggao Liu

## Abstract

Cytotoxic chemotherapy and immune checkpoint inhibitors (ICIs) have transformed the management of advanced cancers, yet durable responses remain restricted to subsets of patients and strongly depend on the tumor immune microenvironment (TIME). Distinct “hot” and “cold” TIMEs differ in pre-existing effector T-cell activity and treatment-induced immunogenicity, suggesting that combination regimens should be tailored to microenvironmental context under realistic clinical constraints. Here we develop a deterministic dynamic optimization framework that jointly designs chemotherapy and ICI dosing schedules on a mechanistic tumor–immune model. The model tracks sensitive and resistant tumor cells, effector CD8^+^ T cells and circulating drug concentrations with pharmacodynamic couplings encoding immunogenic cell death and checkpoint inhibition. We formulate a continuous-time optimal control problem that balances tumor burden against drug exposure while enforcing cumulative dose limits, end-of-interval concentration bounds and bounds on stepwise changes in infusion rates. Using gradient-based optimization with embedded forward sensitivities, we solve the resulting nonlinear programs for four representative TIME phenotypes (extremely cold, hot, cold and cold with high antigenic resistance). The optimizer recovers qualitatively distinct and biologically interpretable regimens, including chemotherapy-dominated schedules in extremely cold TIMEs, aggressive early ICI administration in hot TIMEs and minimal ICI pulses in cold TIMEs. Alternative weightings of treatment objectives reveal trade-offs between tumor eradication and effector preservation. These results show that constrained dynamic optimization can systematically derive phenotype-specific, mechanistically consistent combination strategies and provide a reusable computational module for integrating TIME-resolved models into in silico trial and quantitative systems pharmacology workflows.

**Author summary:** Immune checkpoint inhibitors (ICI) have transformed cancer therapy, yet their benefit varies dramatically across tumor immune microenvironments (TIMEs). In the clinic, ICI is often combined with chemotherapy, and clinicians must decide not only whether to use ICI, but also how to schedule both agents over months of treatment. We develop a reusable dynamic optimization framework that takes any ordinary differential equation model of tumor–immune–therapy interactions as input, enforces clinically motivated dosing and toxicity constraints, and outputs optimal chemo–ICI regimens. Using a mechanistic model and four virtual patients representing hot, cold, and extremely cold TIMEs, we show that the framework automatically suppresses futile ICI dosing in immune deserts, while allocating aggressive combination therapy to highly ICI-responsive tumors.

## 1 Introduction

Cytotoxic chemotherapy and immune checkpoint inhibitors (ICIs) have substantially improved outcomes for many patients with advanced cancer (Topalian et al., 2015). Nevertheless, durable benefit remains confined to a subset of patients and is strongly modulated by the tumor immune microenvironment (TIME), which spans highly inflamed ”hot” tumors to immune-excluded or immune-desert ”cold” phenotypes (Bonaventura et al., 2019; Liu et al., 2019). In triple-negative breast cancer and other aggressive solid tumors, large randomized trials such as KEYNOTE-355 have shown that chemo–ICI combinations can extend progression-free survival compared with chemotherapy alone (Cortes et al., 2020). However, it remains unclear how chemo-ICI regimens should be timed and dosed across distinct TIMEs from hot, inflamed tumors to cold, poorly infiltrated or immune-desert lesions. In practice, scheduling decisions are still guided by empirical protocols rather than mechanism-based, phenotype-specific design (Sharma et al., 2017; Ribba et al., 2018; El Haout et al., 2021).

Mathematical models of tumor-immune dynamics offer a principled way to explore such design questions. Early work by de Pillis and colleagues formulated coupled ordinary differential equation (ODE) models for tumor cells, immune effector cells, and cytotoxic drugs, and analyzed mixed chemo-immunotherapy strategies together with their biological interpretation (de Pillis et al., 2006; Castiglione and Piccoli, 2007). Subsequent studies have systematically reviewed and extended this modeling paradigm to capture treatment resistance, spatial heterogeneity, and diverse features of the TIME (Yin et al., 2019; Wang and Schättler, 2016; Das et al., 2021; Padmanabhan et al., 2020). More recent models have focused on specific indications and modalities, including androgen-deprivation plus vaccine and ICI therapy in metastatic castration-resistant prostate cancer, radiotherapy combined with CTLA-4 blockade (Kim et al., 2023; Ledzewicz and Schättler, 2024; Ledzewicz et al., 2012), and PD-1/PD-L1 inhibitors in advanced melanoma and other solid tumors (Butner et al., 2020, 2021; Liu et al., 2019; Khalili and Vatankhah, 2023; Sardar et al., 2024; Nave, 2022; Santra and Samanta, 2025). In parallel, high-throughput molecular profiling and deconvolution tools such as CIBERSORT have enabled detailed quantification of tumor-infiltrating immune populations, providing rich data to inform and constrain mechanistic TIME models (Azizi, 2025; Najafi and George, 2025).

Although these studies underscore the value of mathematical oncology, two methodological gaps remain. First, most existing works either explore fixed, pre-specified regimens or perform limited parametric sweeps over dosing schedules (Ledzewicz et al., 2024; Omer et al., 2025; Baker et al., 2025; Dodek et al., 2025). As a result, they do not fully exploit the available control degrees of freedom, especially when therapies can in principle be titrated continuously in time, and they rarely encode clinical constraints on cumulative dose and immune-related toxicity in a systematic way. Second, while recent deep reinforcement learning (RL) approaches have been applied to continuous cancer therapy optimization (Andersona et al., 2024; Qin and Wu, 2025), such controllers typically require extensive simulation or clinical data for training, may converge to policies that are difficult to interpret mechanistically, and can be challenging to verify or tune in safety-critical applications. These considerations leave open whether one can systematically compute phenotype-adaptive, clinically constrained regimens using transparent dynamical models, without requiring large-scale clinical data or opaque learned policies.

Here we develop a deterministic dynamic optimization framework for chemo–ICI scheduling across heterogeneous TIMEs. First, we formulate a six-dimensional ODE model of sensitive and resistant tumor cells, effector CD8^+^ T cells, and circulating ICI and chemotherapy, calibrated to represent four canonical TIME phenotypes: extremely cold, hot, cold, and cold with high resistant–cell growth. Second, we embed this model in a constrained optimal-control framework that enforces cumulative dose limits, pharmacokinetic safety caps, and ramp constraints on dose changes, and we solve for continuous-time dosing profiles using gradient-based nonlinear programming with embedded forward sensitivities. Third, by applying this reusable framework to four virtual TIME scenarios under two clinical preference settings, we show that the optimizer autonomously suppresses futile ICI in cold TIMEs, saturates synergistic chemo–ICI in hot TIMEs, and reveals trade-offs between tumor debulking and long-term effector preservation, thereby generating testable hypotheses for phenotype-stratified trial design. Unlike prior optimal-control studies that either assume idealized dosing patterns or focus on a single tumor context, we develop a reusable dynamic optimization module that (i) enforces clinically motivated dosing and toxicity constraints, and (ii) systematically compares phenotype-specific chemo-ICI regimens across distinct TIME settings.

## 2 Tumor–immune dynamical model

We adopt a mechanistic ordinary differential equation model to describe the coupled dynamics of tumor cells, effector immune cells, and two systemic therapies: an immune checkpoint inhibitor (ICI) and a cytotoxic chemotherapeutic agent. The model builds on previous work on tumor–immune interactions and mixed chemo–immunotherapy (de Pillis et al., 2006; Yin et al., 2019; Butner et al., 2020, 2021; Kim et al., 2023), and on recent formulations of hot versus cold tumor immune microenvironments. The schematic TIME landscape under ICI and chemotherapy is illustrated in Fig. 1. In this section we summarize the state variables, control inputs and interaction terms that define the dynamical system; the subsequent section embeds this model in a constrained optimal control framework.

**Figure 1:**
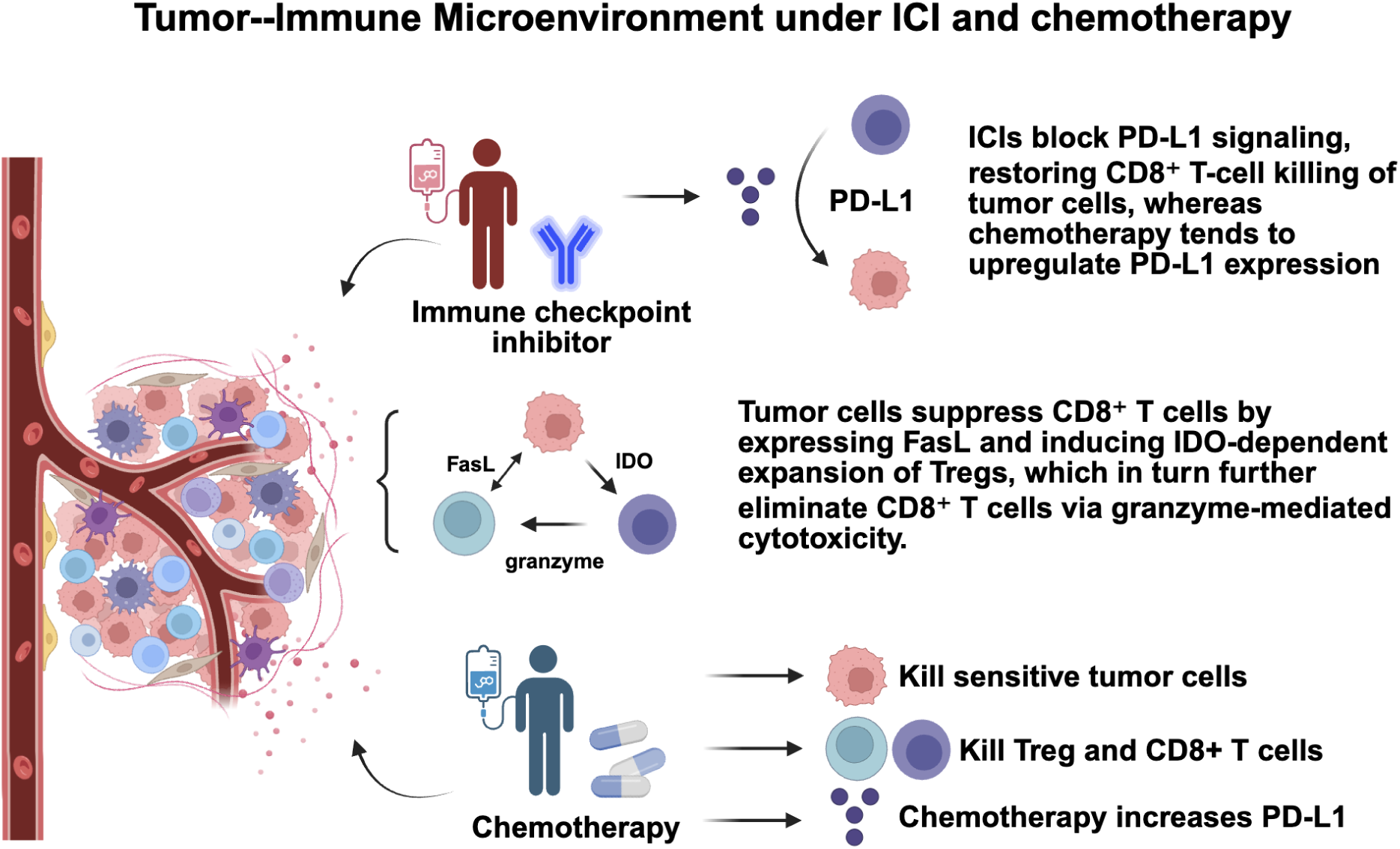
Tumor–immune microenvironment under ICI and chemotherapy. Created in https://BioRender.com.

### 2.1 State variables and control inputs

We built a parsimonious ODE model that tracks (i) sensitive and resistant tumor cells, (ii) effector CD8^+^ T cells, and (iii) the pharmacokinetics and pharmacodynamics of an ICI and a cytotoxic drug. The model captures three key processes: logistic competition between sensitive and resistant tumor subpopulations, immune-mediated killing modulated by checkpoint inhibition, and chemo-induced damage to both tumor and effector cells. Distinct TIME phenotypes are encoded by varying parameters that control baseline immune infiltration, checkpoint strength, and chemo-induced toxicity. We consider five primary state variables as follows.

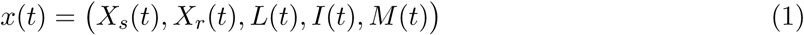

where *X_s_*(*t*) denotes the burden of treatment-sensitive tumor cells, *X_r_*(*t*) denotes the burden of resistant tumor cells, *L*(*t*) denotes effector CD8^+^ T cells in the tumor or peripheral compartment, *I*(*t*) denotes the concentration of ICI, and *M* (*t*) denotes the concentration of chemotherapy. The total tumor burden is defined as

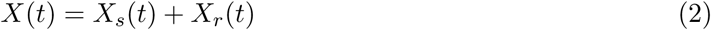

Two time-dependent control inputs represent externally administered therapies:

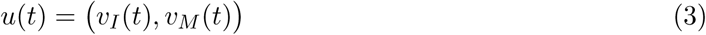

where *v_I_*(*t*) and *v_M_* (*t*) denote the effective dosing rates of ICI and chemotherapy, respectively. These controls are subject to bounds and cumulative dose constraints, as detailed in the dynamic optimization framework below.

### 2.2 Tumor growth, immune activation, and drug effects

The tumor components obey logistic growth with competition between sensitive and resistant sub-populations. Denoting by *X*_max_ the effective carrying capacity and by *α_sr_, α_rs_* the cross-competition coefficients, the net growth rates of *X_s_* and *X_r_* are modulated by immune-mediated killing and chemo-induced damage. The full ODE system is described as follows:

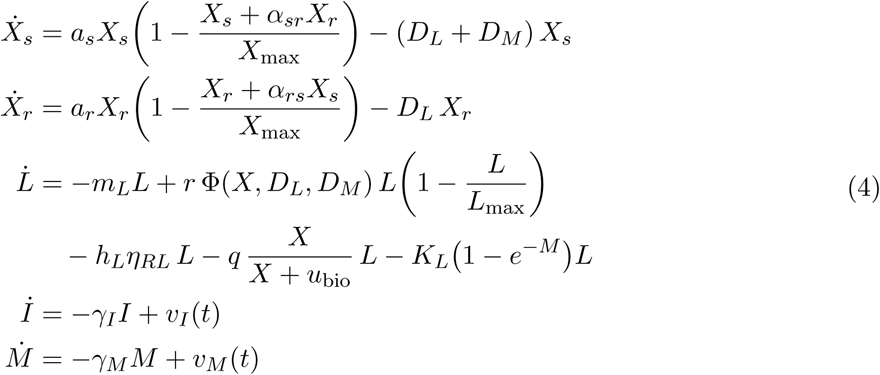

where *a_s_* and *a_r_* are intrinsic proliferation rates of sensitive and resistant cells, *m_L_* is the natural death rate of effector T cells, *L*_max_ is the effective carrying capacity of *L*, and *γ_I_, γ_M_* are first-order clearance rates of the ICI and chemotherapy, respectively. The terms multiplied by *h_L_*, *q* and *K_L_* capture suppression of T cells by regulatory mechanisms, tumor burden and chemo-induced toxicity.

Effector T cell stimulation is modeled through a composite signal

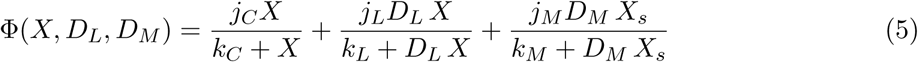

where the first term represents antigen presentation due to baseline immunogenic cell death, the second term reflects immunogenic cell death enhanced by CD8^+^ T cells, and the third term accounts for chemo-induced immunogenicity in the sensitive tumor compartment.

The quantities *D_L_*(*t*) and *D_M_* (*t*) encode, respectively, effective immune-mediated killing and chemo-induced killing of sensitive tumor cells. They depend on the instantaneous checkpoint inhibition level *µ*(*t*) and chemotherapy exposure:

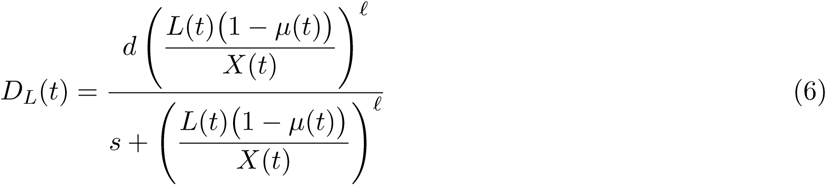

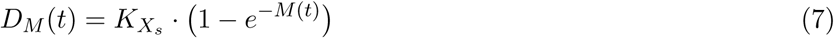

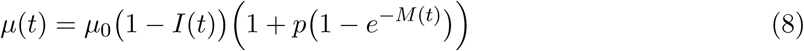

Here *d*, *s* and *ℓ* shape the saturation of immune-mediated killing as a function of the effector-to-tumor ratio, *µ*_0_ represents the baseline strength of checkpoint-mediated T cell suppression in the absence of treatment, and *p* captures chemo-induced up-regulation of inhibitory ligands on tumor or immune cells. The parameter *K_X_s* scales chemo-induced killing in the sensitive compartment.

Equations (6)–(8) compactly encode the influence of the TIME phenotype on treatment response. “Hot” tumors with pre-existing immune infiltration and weaker checkpoint suppression correspond to lower *µ*_0_ and/or higher *j_L_*, whereas “cold” or highly immunosuppressive TIMEs exhibit larger *µ*_0_, stronger tumor-mediated suppression *q* and more pronounced chemo toxicity to T cells (larger *K_L_*). In the section of case studies, we specify representative parameter sets for extremely cold, hot, cold and cold-with-high-resistance-growth phenotypes and analyze the corresponding optimized treatment strategies.

## 3 Dynamic optimization framework

Building on the tumor–immune dynamical system, we formulate the joint chemotherapy and ICI scheduling task as a continuous-time dynamic optimization problem. The core idea is to parameterize the dosing schedule as a finite set of per-interval infusion rates, embed biologically motivated toxicity and smoothness constraints, and then solve the resulting finite-dimensional nonlinear programming using gradient-based methods. In this section we describe the mathematical structure of the dynamic optimization framework (DOF), while the numerical implementation details are provided in next section. Although we instantiate the framework on a specific chemo–ICI TIME model, the DOF is model-agnostic and can in principle be applied to any ODE-based tumor–immune system with suitable state and control definitions.

### 3.1 Time horizon and treatment phases

The simulated clinical trajectory consists of three phases: a pre-treatment period (tumor and immune dynamics without therapy), an active treatment phase (optimized combination of ICI and chemotherapy), and a post-treatment follow-up (drug washout and natural evolution). In the optimization problem, only the active treatment phase is controlled explicitly.

Let [*t*_0_*, t_f_*] denote the treatment horizon in *weeks* (e.g. *t*_0_ = 0, *t_f_* = 30 in the numerical experiments), and let *x*^(*t*) = (*X* (*t*)*, X* (*t*)*, L*(*t*)*, I*(*t*)*, M* (*t*)*, J* (*t*))⊤ denote the 6-dimensional augmented state, where *J* (*t*) accumulates the running cost. The controlled dynamics over [*t*_0_*, t_f_*] are given by

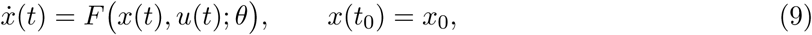

where *u*(*t*) = (*v_I_*(*t*)*, v_M_* (*t*))^⊤^ is the control vector and *θ* collects all fixed model parameters.

### 3.2 Control vector parametrization and pulsed dosing

To obtain a finite-dimensional decision space, we adopt a control vector parametrization (CVP). The treatment horizon [*t*_0_*, t_f_*] is partitioned into *N* equal sub-intervals [*t_k_, t_k_*_+1_)*, k* = 0*, …, N* − 1. On each sub-interval [*t_k_, t_k_*_+1_) we introduce constant nominal infusion rates *u_k_* = (*v_I,k_, v_M,k_*)*, k* = 0*, …, N* − 1 and collect all decision variables into *z* = (*v_I,_*_0_*, v_M,_*_0_*, …, v_I,N_*_−1_*, v_M,N_*_−1_)⊤ ∈ R^2^*^N^*. In order to emulate clinically realistic bolus-like administration while preserving the same total dose, each sub-interval is further split in the ODE integration into a short pulse subsegment followed by a washout subsegment. Concretely, we define:

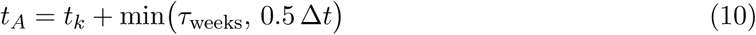

where *τ*_weeks_ is the prescribed pulse duration (e.g. three days expressed in weeks) and Δ*t* = *t_k_*_+1_−*t_k_*. On the pulse subsegment [*t_k_, t_A_*) we apply a scaled rate

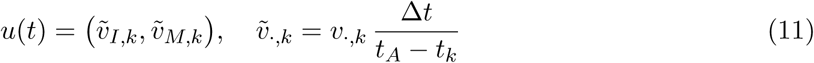

so that the area under the curve satisfies

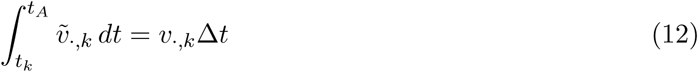

This construction ensures that the total integrated dose over each interval matches the nominal CVP parameters while yielding bolus-like input profiles.

### 3.3 Objective functional and clinical trade-offs

Rather than computing the cost via explicit quadrature outside the ODE solver, we represent the running cost as an additional state variable *J* (*t*) with *J* (*t*_0_) = 0. The cost functional is not intended to match a specific clinical endpoint, but rather to encode a tunable trade-off between tumor burden, drug use, and preservation of effector T cells, which we explore systematically in the numerical experiments. We define

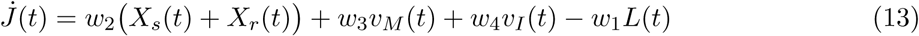

where *w*_1_*, w*_2_*, w*_3_*, w*_4_ *>* 0 are scalar weights that encode the trade-off between tumor burden, effector T-cell preservation, and drug usage, and can be interpreted as reflecting different clinical preferences. The continuous-time performance index is then simply the terminal value J (*z*) = *J* (*t_f_* ; *z*):

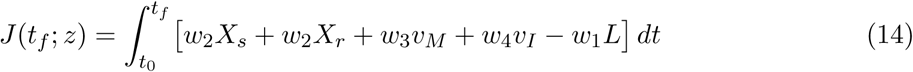

which in the implementation is read directly as the sixth state component at *t_f_*. This construction avoids numerical inconsistencies between the ODE solver and a separate quadrature routine and makes sensitivity computation straightforward.

### 3.4 Clinical constraints on dose, exposure and ramps

The admissible control set is characterized by three families of constraints that reflect clinical and numerical considerations.

#### (i) Per-cycle dose limits and treatment windows

Each per-interval decision variable is subject to lower and upper bounds:

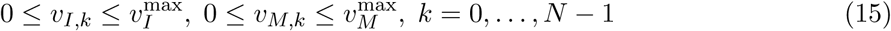

In addition, chemotherapy and ICI are restricted to predefined treatment windows. Let *K_I_* and *K_M_* denote index sets of intervals where ICI and chemotherapy are allowed, respectively. Outside these windows the corresponding upper bounds are set to zero in the code, effectively forcing *v_I,k_* = 0 or *v_M,k_* = 0.

#### (ii) Cumulative dose caps

Total exposure of ICI and chemotherapy is limited via

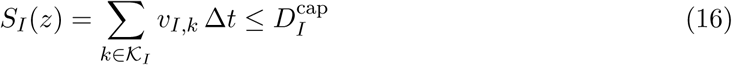

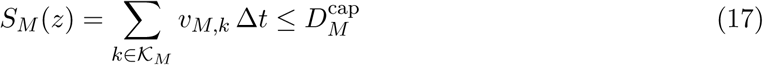

where 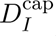 and 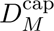 are specified in week units in the implementation (reflecting the total integrated dose). These constraints are encoded as a following smooth inequality map with an analytic Jacobian.

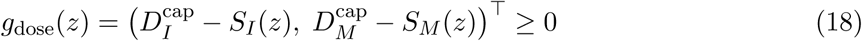

#### (iii) Pharmacokinetic safety caps

To prevent unphysiologically high systemic drug levels we impose upper bounds on ICI and chemotherapy concentrations evaluated at the end of each sub-interval. Let (*I_k_, M_k_*) denote the values of (*I, M*) at *t_k_*_+1_ obtained from forward integration with a given *z*. Then

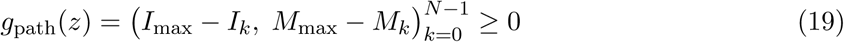

where *I*_max_ and *M*_max_ are prescribed upper bounds. In the present work we deliberately relax lower bounds on *L*(*t*) and more detailed toxicity metrics in order to keep the optimization numerically well-conditioned.

#### (iv) Ramp constraints on dose changes

Finally, to avoid unrealistic step-to-step fluctuations in dosing, ramp constraints limit the change between neighboring decision variables:

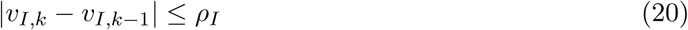

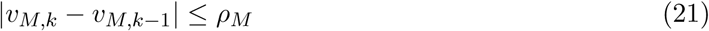

for *k* = 1*, …, N* − 1 and prescribed slopes *ρ_I_, ρ_M_ >* 0. In the implementation these are represented as linear inequality functions *g*_ramp_(*z*) ≥ 0 with an explicit sparse Jacobian.

### 3.5 Numerical implementation: single-shooting NLP with embedded sensitivities

Given the dynamic optimization formulation above, we discretize the control inputs by control vector parametrization and solve the resulting finite-dimensional nonlinear program in a single-shooting fashion. In this approach, all decision variables are the per-interval dosing parameters, while the state trajectory is obtained purely by numerical integration of the ODE system.

For a candidate control vector:

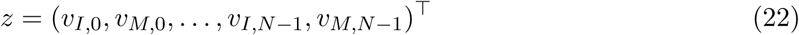

we reconstruct a piecewise-constant, pulsed control *u*(*t*) over the treatment horizon. The augmented state

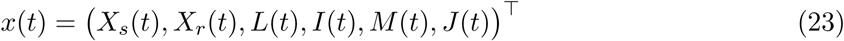

is then propagated from *t*_0_ to *t_f_* under this control by integrating the tumor–immune–therapy ODE system. The performance index is read directly as the sixth component at final time, *J* (*z*) = *J* (*t_f_* ; *z*), because the running cost is accumulated through *J̇*(*t*).

To provide gradients to the NLP solver, we embed forward sensitivity equations inside the ODE integration. Let *P* = 2*N* denote the number of control parameters and

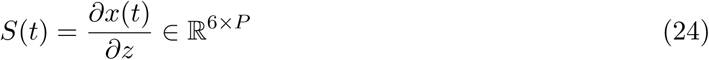

collect the sensitivities of the state with respect to an internal parameter vector *z* that stores all per-interval doses. The sensitivity dynamics follow the standard linearized form

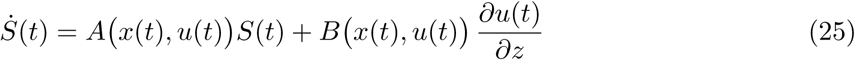

where *A* = *∂F/∂x* and *B* = *∂F/∂u* are the Jacobians of the right-hand side *F* of the ODE system with respect to the state and control variables. Within each treatment interval, the mapping *∂u/∂z* is realized by sparse “injection” matrices that pick out the two active parameters (*v_I,k_, v_M,k_*); outside the active intervals this mapping is zero. For numerical integration, we concatenate *x*(*t*) and vec(*S*(*t*)) into a single augmented state and integrate the combined system using an adaptive ODE solver.

Because the running cost has been absorbed into *J* (*t*), the gradient of the objective with respect to the internal parameters is obtained simply from the sensitivities at final time:

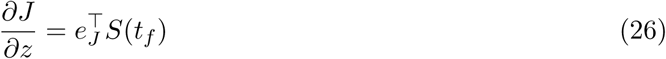

where *e_J_* selects the *J* -component of the state. The optimizer operates on a decision vector *z* that interleaves ICI and chemotherapy doses; a fixed permutation matrix is used to map between the internal parameter ordering and *z*, yielding the gradient *∂J/∂z* reported to the solver. The same sensitivity information is also used to assemble Jacobians of inequality constraints that depend on the state, such as the pharmacokinetic safety caps on *I* and *M* at the end of each treatment interval. Full analytical expressions for the Jacobian matrices and sensitivity equations are provided in Supplementary Text.

The resulting nonlinear programming problem, equipped with the analytic gradients and constraint Jacobian, is solved using the trust-constr method. The Hessian is approximated by a quasi-Newton update; the user-specified tolerances are *gtol* = 10^−6^, *xtol* = 10^−8^, *maxiter* = 200.

If the trust-region solver fails to converge or yields an infeasible point (negative minimum inequality residual), the current iterate is passed as an initial guess to SLSQP as a robust fallback. This two-stage strategy, combined with embedded sensitivities and the LSODA integrator, yields numerically stable and reproducible optimal regimens for all cold and hot tumor scenarios considered in following Section.

## 4 Results on phenotype-dependent optimal regimens across tumor immune microenvironments

To illustrate the behavior of the proposed dynamic optimization framework and to compare it with TIME-based insights in the literature, we conduct two sets of numerical experiments. In both sets, the tumor–immune ODE parameters are chosen such that the four baseline scenarios correspond to the canonical TIME phenotypes used in the reference study (extremely cold, hot, cold and cold with high resistant growth). What differs between the two experiments is how the decision-level hyper-parameters in the DOF (trade-off weights and cumulative dose caps) are specified:

- Case 1: a uniform set of cost weights (*w*_1_*, w*_2_*, w*_3_*, w*_4_) and dose caps 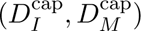 is used for all four TIME scenarios; only the biological parameters defining the TIME phenotypes differ.
- Case 2: the underlying TIME parameters are kept identical to Experiment 1, but the DOF hyper-parameters are adjusted to match the case-specific settings in the reference work, yielding more ICI-favoring caps and weights in hot TIME and more chemo-oriented caps in extremely cold TIME.

In both experiments the treatment horizon is same, with a pre-treatment period of approximately 400 days and a post-treatment follow-up up to about 700 days. The control vector parametrization uses *N* = 10 intervals, with ICI and chemotherapy restricted to the predefined windows K*_I_* and K*_M_* as described in Section 3. For each scenario the NLP is solved using the trust-constr method with analytic gradients and a fallback to SLSQP when needed.

### 4.1 Phenotype heterogeneity alone yields distinct optimal chemo-ICI strategies (case 1)

To isolate the impact of TIME heterogeneity on optimal treatment design, we first simulated four virtual patients under an identical set of pharmacodynamic and control parameters: (a) an extremely cold TIME, (b) a hot TIME, (c) a moderately cold TIME, and (d) a cold TIME with elevated resistant-cell proliferation *a_r_*. All models shared the same toxicity coefficients, maximum dosing limits, and temporal segmentation (Δ*t* = 3 weeks); only the initial immune infiltration and intrinsic proliferation rates differed across cases. The resulting tumor trajectories, immune-cell dynamics, and control profiles are summarized in Fig. 2, with quantitative outcome metrics in Table. 2. Together, these simulations ask a single question: how far can phenotype heterogeneity alone drive divergence in the “best” chemo-ICI strategy when clinical constraints are held fixed?

**Figure 2:**
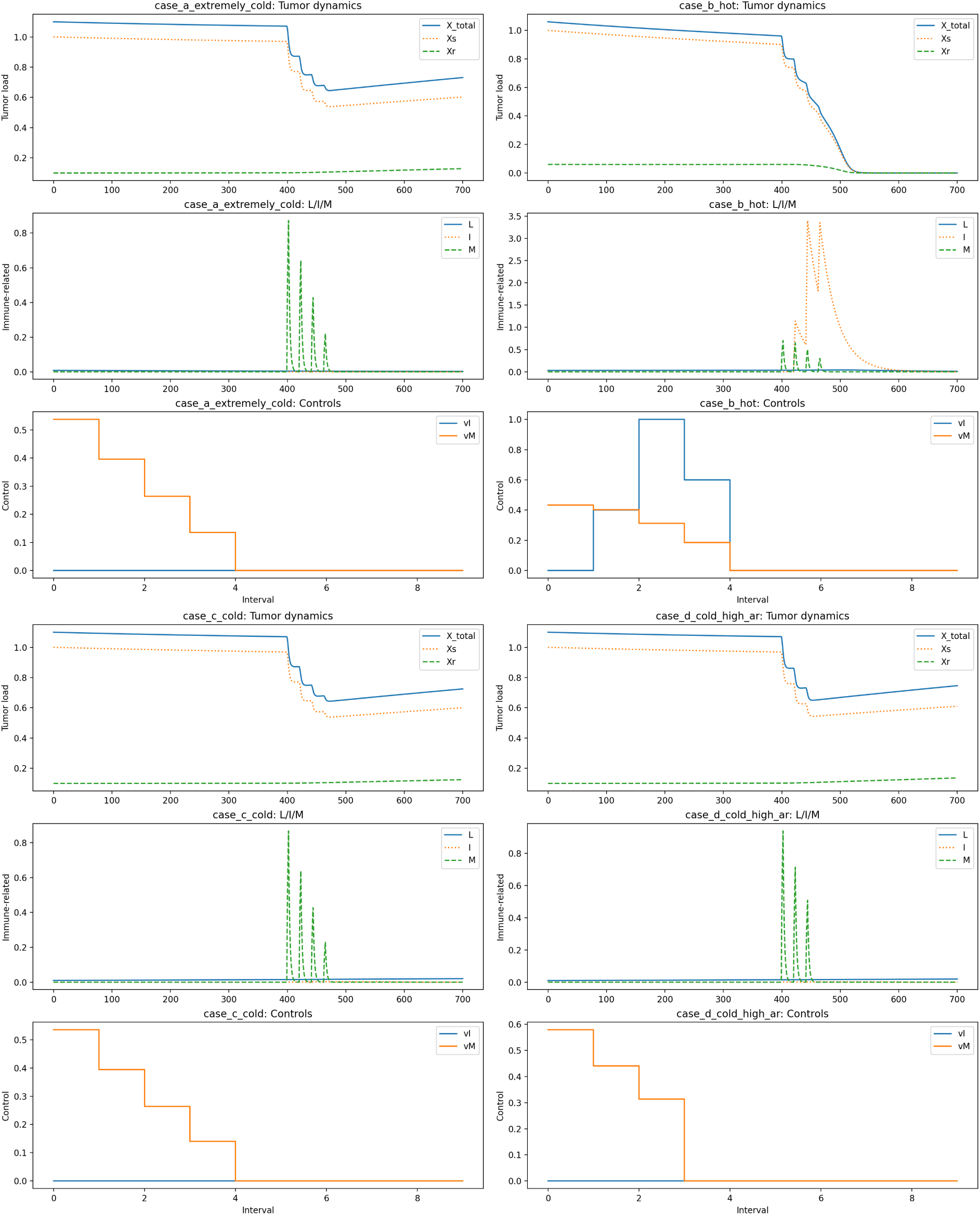
Overview of tumor, immune, and control dynamics under identical parameters for four TIME phenotypes. Each sub-panel shows the time evolution of tumor burden (*X*_total_*, X_s_, X_r_*), immune-related variables (*L, I, M*), and optimal control inputs (*v_I_, v_M_*).

#### (i) Tumor regression dynamics

Among the four phenotypes, the hot TIME (case B) achieves complete tumor clearance, with the total tumor load *X*_total_ dropping from 1.02 to nearly zero between days 420 and 500 (Table 2 and Fig. 3). In contrast, all three cold phenotypes retain a non-zero residual burden at the end of follow-up (*X*(*t_f_*) ∈ [0.73, 0.75]), despite receiving essentially identical cumulative chemotherapy exposure (*S_M_* ≈ 28). The minimum tumor burden in these cold-type TIMEs is *X*_min_ ≈ 0.65, corresponding to only a roughly 35% transient shrinkage before regrowth.

**Figure 3:**
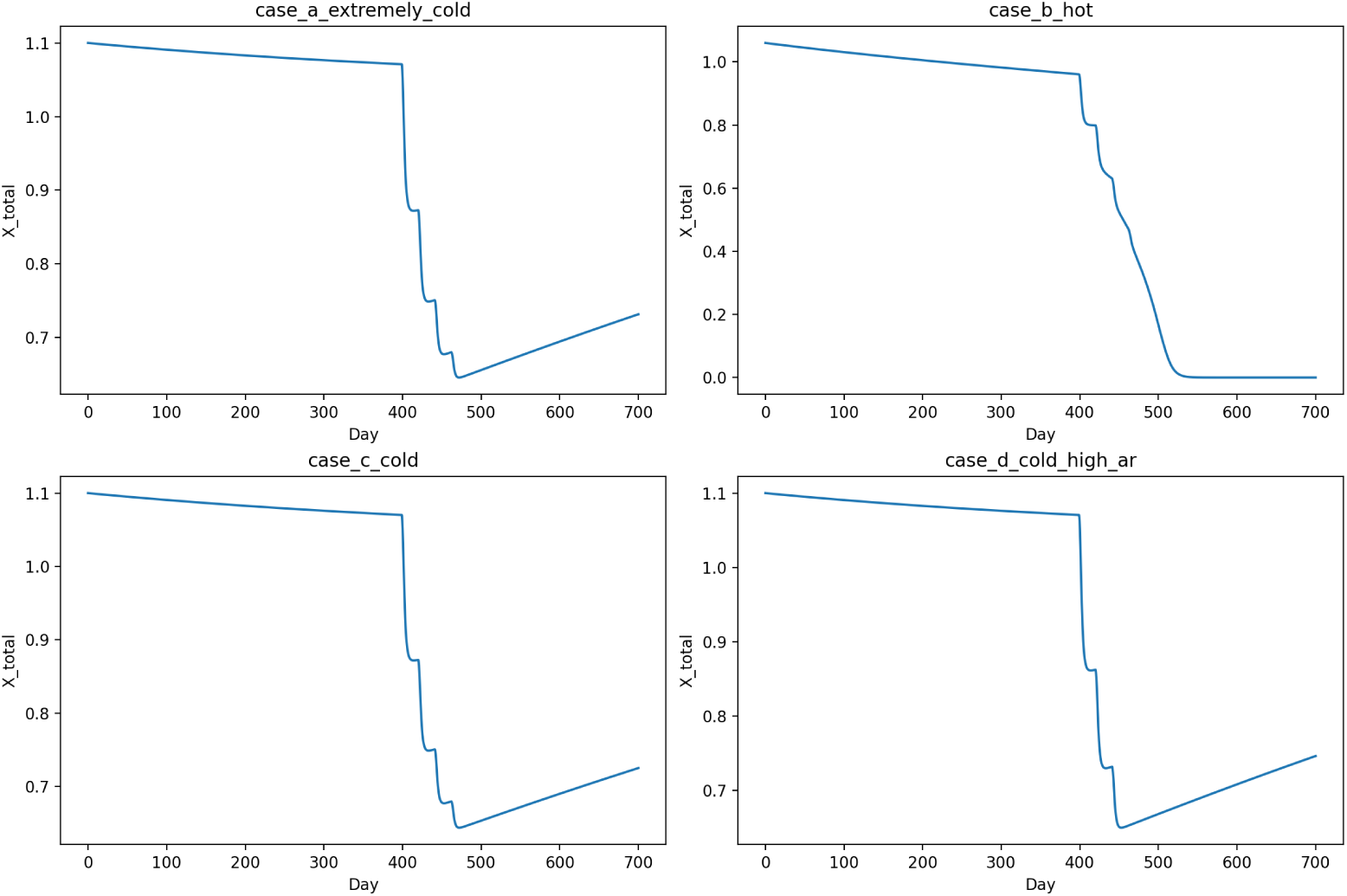
Comparison of total tumor burden trajectories *X*_total_(*t*) under identical parameter settings. Each curve corresponds to one TIME phenotype (A–D), showing distinct regression and relapse patterns.

Figure 3 directly compares *X*_total_(*t*) across the four cases under identical pharmacodynamic parameters. The hot phenotype (case B) exhibits a monotonic, nearly exponential decay leading to eradication around *t* ≈ 480 days. By contrast, the extremely cold and cold tumors (cases A, C, and D) show a biphasic pattern: an initial partial response followed by relapse once cytotoxic pressure relaxes. Quantitatively, the separation between *X*_min_ ≈ 0.65 for cold-type TIMEs and *X*_min_ ≈ 0 for the hot TIME underscores that pre-existing immune activity, rather than total chemotherapy exposure, is the primary determinant of durable tumor control.

#### (ii) Immune-cell activation and control allocation

Figure 4 displays the effector lymphocyte trajectories *L*(*t*) for the four TIMEs. In the hot phenotype, *L*(*t*) undergoes a pronounced amplification following the second ICI pulse, reaching *L*_max_ = 0.042 and closely tracking the regression of *X*_total_. In all cold phenotypes, by contrast, effector responses are weak: *L*(*t*) either remains near its baseline value or gradually decays due to insufficient antigenic stimulation, with *L*_max_ *<* 0.021. This pattern quantitatively mirrors immune exclusion in non-inflamed tumors, as reported experimentally for “cold” microenvironments Bonaventura et al. (2019).

**Figure 4:**
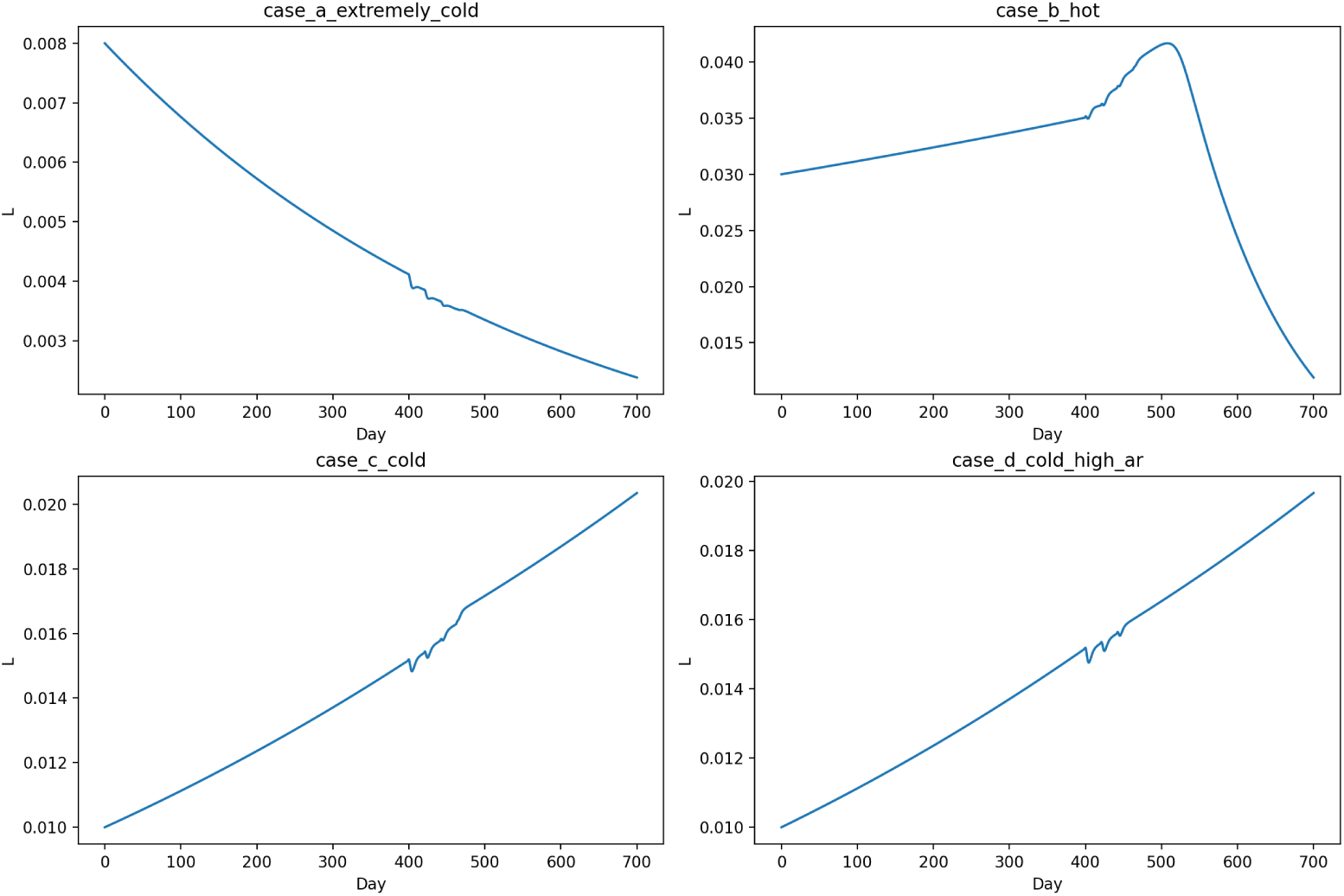
Comparative dynamics of effector lymphocytes *L*(*t*) under identical pharmacodynamic parameters across four TIME phenotypes (A–D). The hot phenotype sustains higher *L*(*t*) amplitudes and delayed decay, whereas cold and extremely cold cases exhibit weak activation and early immune exhaustion.

The optimal control schedules reflect this dichotomy. In the extremely cold TIME (case A), the optimizer allocates no ICI at all (*S_I_* = 0), effectively recognizing that PD-1/PD-L1 blockade provides negligible marginal benefit when effector precursors are scarce. In case B (hot TIME), both the ICI and chemotherapy budgets are fully exploited (*S_I_* = 42, *S_M_* = 28), with synchronized pulses that yield complete remission. The moderately cold and high-*a_r_* cases receive only small compensatory ICI doses (*S_I_ <* 0.12), with chemotherapy acting as the primary driver of tumor debulking. Thus, even under identical constraints, the optimizer redistributes control effort in a phenotype-specific way: intensive combined chemo-ICI for hot TIMEs and chemotherapy-dominated regimens for cold TIMEs.

#### (iii) Mechanistic interpretation

These differences arise from how TIME parameters modulate the local sensitivities of tumor growth to immune and drug effects. In the hot TIME, strong coupling between ICI concentration and tumor lysis (large negative *∂f_X_s /∂I*) yields steep gradients of the objective with respect to ICI dosing. The optimizer therefore “sees” large payoffs from ICI administration and drives the solution toward near-maximal immunotherapy use. In cold microenvironments, the same derivatives are much smaller, reflecting weak immune-mediated killing; consequently, gradients with respect to ICI are suppressed and the algorithm shifts the optimal solution toward purely chemotherapeutic schedules. In this sense, the dynamic optimization framework learns an implicit and clinically intuitive policy: ICI is deployed aggressively only when sufficient immune infiltration exists to support it.

#### (iv) Summary of identical-parameter experiments

Under identical pharmacodynamic settings, the proposed framework produces internally consistent yet phenotype-specific schedules that exploit the underlying TIME. Hot tumors benefit from high-intensity, temporally coordinated chemo-ICI therapy and achieve full remission; cold and extremely cold tumors remain largely refractory, exhibiting only partial shrinkage followed by regrowth. The automatic suppression of futile ICI dosing in “immune deserts” indicates that the optimizer internalizes a mechanistic hierarchy between immune responsiveness and drug toxicity—a property that is difficult to encode in fixed, hand-crafted regimens.

### 4.2 Clinical preference weighting exposes trade-offs between tumor debulking and effector preservation (case 2)

We next examined how clinical preference weighting and dose caps shape the optimal regimens. To this end, we repeated the four virtual-patient experiments under an alternative parameterization of the objective function and cumulative dose limits, while keeping all TIME parameters identical to case 1 (Table. 1). The resulting trajectories are summarized in Fig. 7, with cross-case comparisons of effector dynamics and tumor burden shown in Figs. 5–6 and quantitative metrics reported in Tables. 3 and 4. In this setting, the question is no longer “which phenotype responds best,” but rather how the optimizer trades tumor debulking against long-term effector preservation when the relative weights on drug cost and immune benefit are altered.

**Figure 5:**
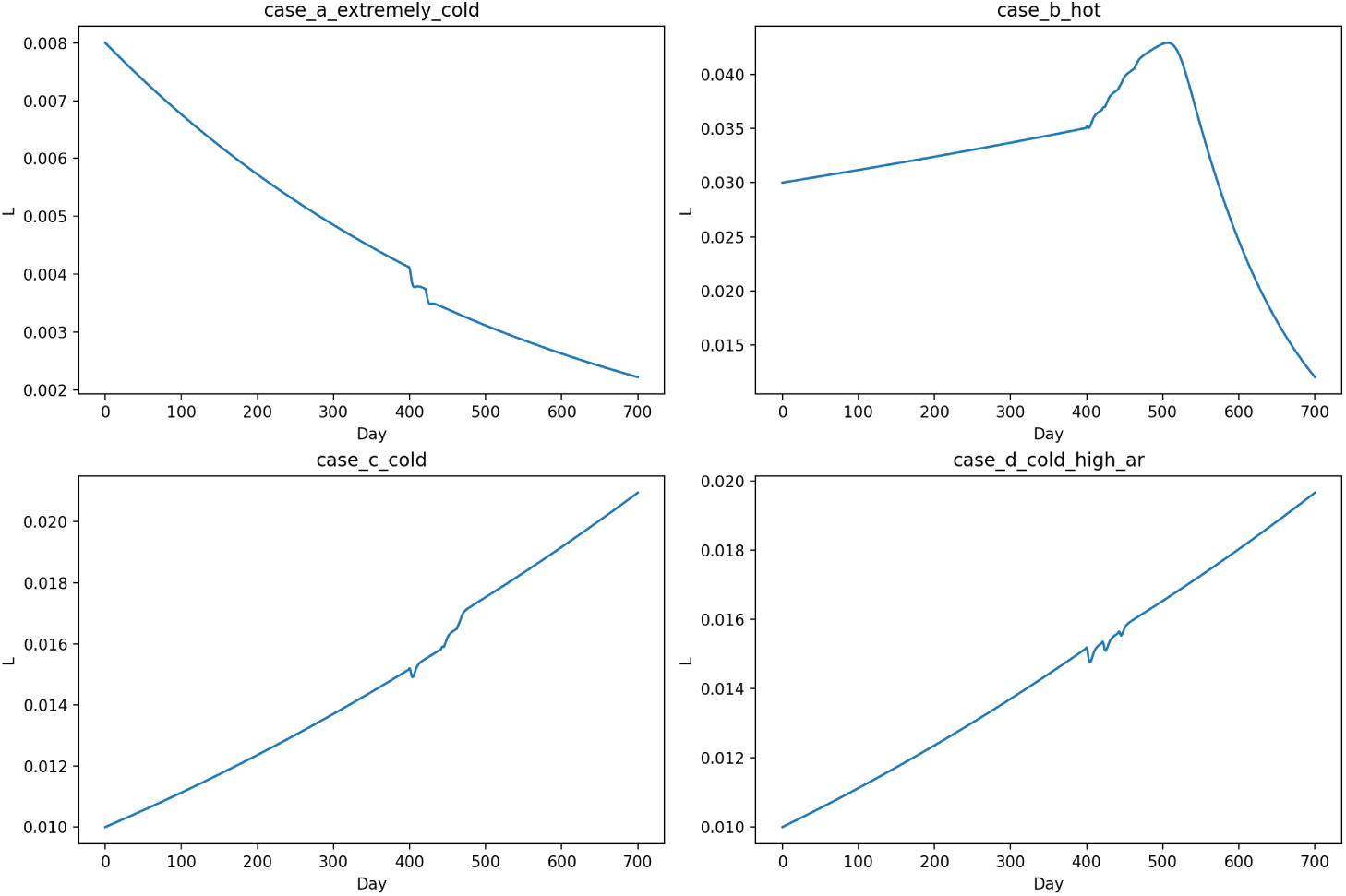
Time evolution of effector lymphocyte density *L*(*t*) under different parameterizations. The four trajectories correspond respectively to: (a) extremely cold, (b) hot, (c) cold, and (d) cold with high antigenic resistance cases. The alternative parameterization reveals distinct immune activation patterns— notably a transient amplification under the hot case and a monotonic decline in the extremely cold case.

**Figure 6:**
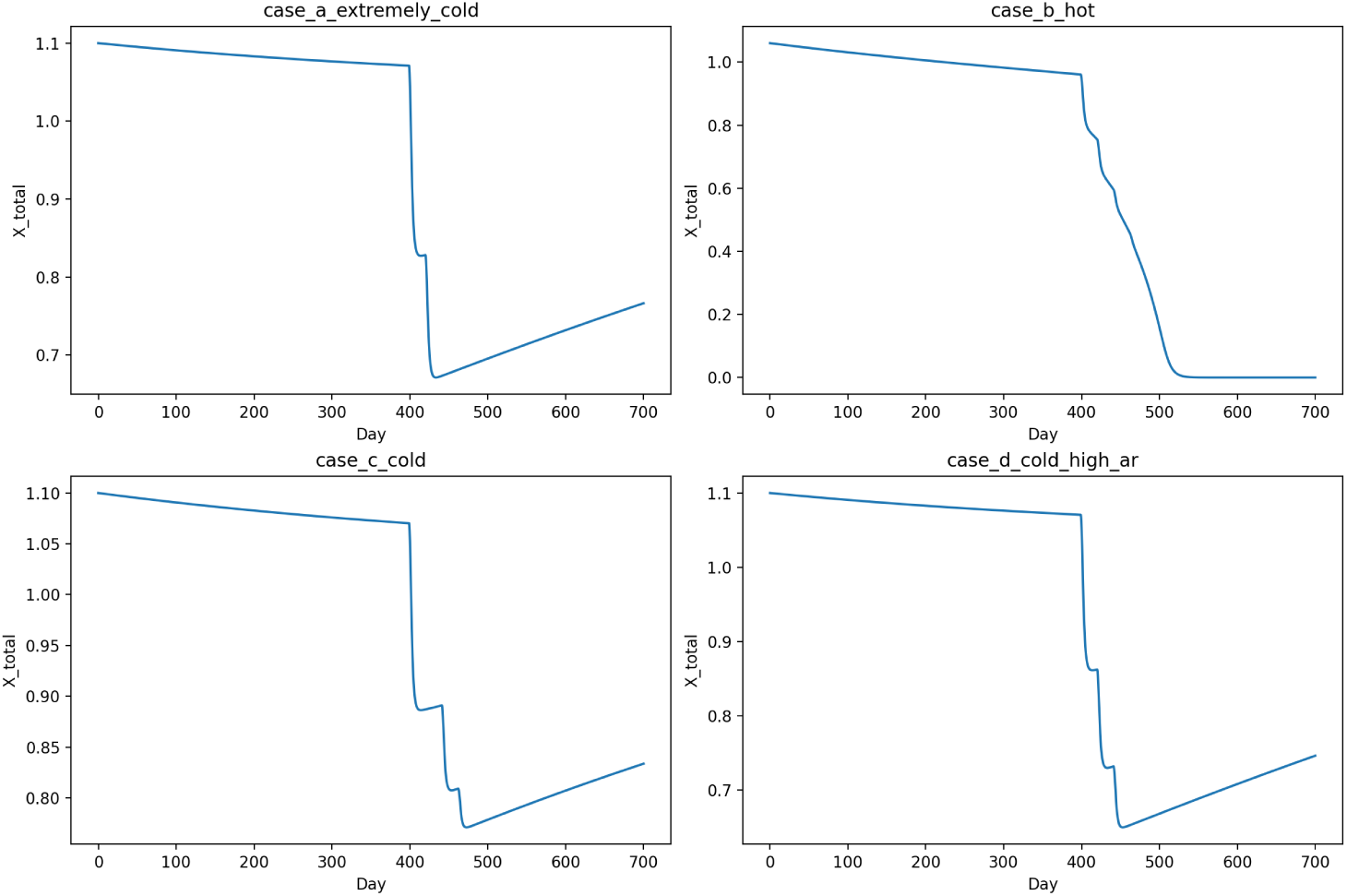
Time evolution of the total tumor burden *X*_total_(*t*) under different parameterizations. The four trajectories correspond respectively to: (a) extremely cold, (b) hot, (c) cold, and (d) cold with high antigenic resistance cases. The parameter variations yield markedly different tumor suppression outcomes— full eradication in the hot scenario, partial regression with rebound in the cold and cold–high antigenic resistance cases, and only transient reduction in the extremely cold case.

**Figure 7:**
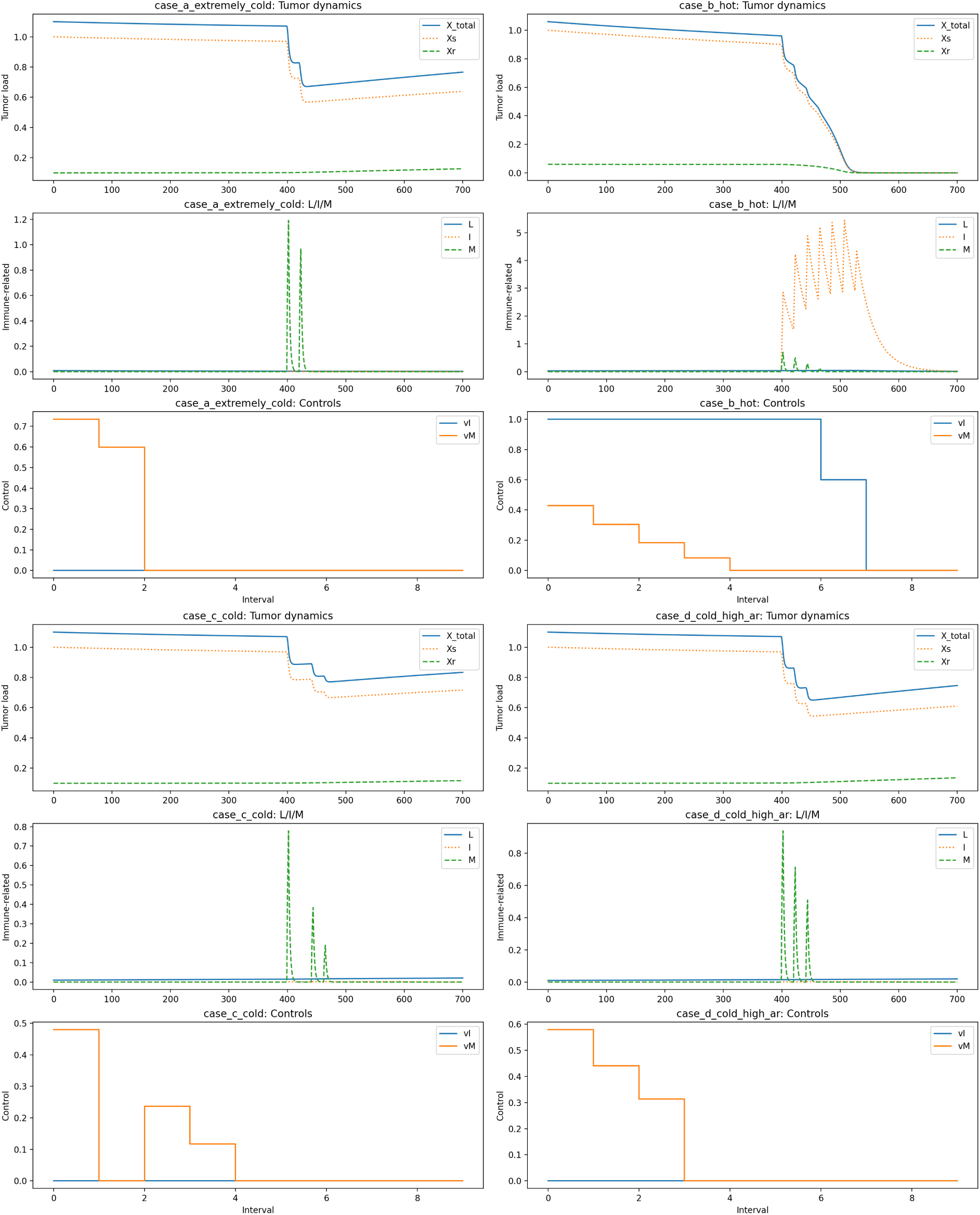
Overview of tumor and immune dynamics under different parameter settings. The four subfigures correspond to the alternative parameterization cases: (a) extremely cold, (b) hot, (c) cold, and (d) cold with high antigenic resistance. Each panel shows the trajectories of tumor burden, effector lymphocytes, and treatment dosing schedules over the simulated period.

**Table 1:**
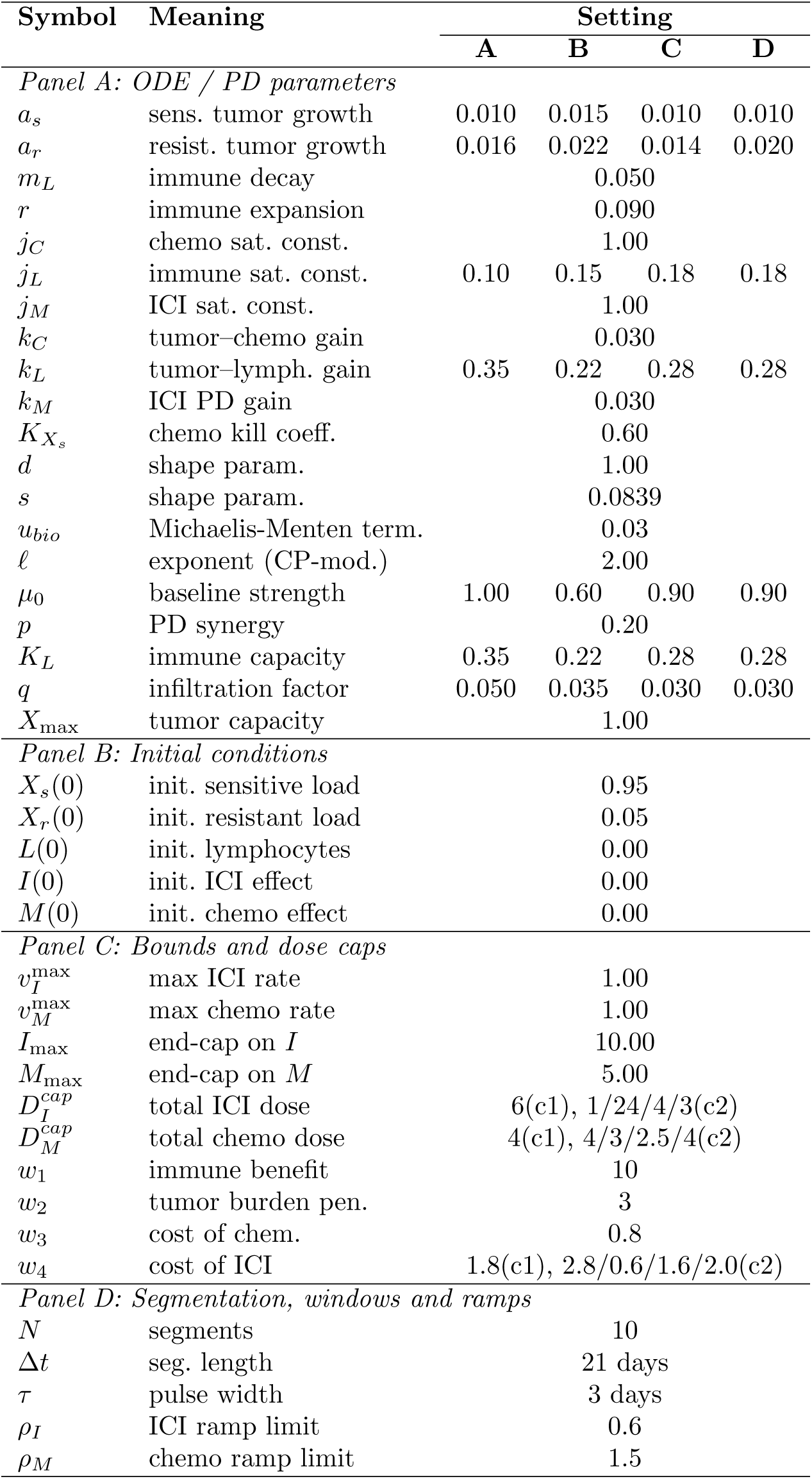
Parameters, initial conditions, bounds, windows and solver settings for two cases (Settings A–D). Columns (A)–(D) correspond to the four TIME scenarios: (A) extremely cold; (B) hot; (C) cold; (D) cold with high resistant growth. C1: case1 and C2: case2.

**Table 2:** Quantitative metrics of tumor and immune dynamics under identical parameter settings. *X*_min_ denotes the minimum total tumor burden *X_s_* + *X_r_*observed during the active treatment window (days 399–609), *X*(*t_f_*) is the final tumor burden at *t_f_*= 700 days, *S_I_* and *S_M_* are the cumulative (time–integrated) doses of ICI and chemotherapy over the 10 treatment intervals, and *L*_max_ is the peak effector lymphocyte level over the whole simulation horizon. All values are dimensionless and normalized.

| Case | $X_{\min}$ | $X(t_f)$ | $S_I$ | $S_M$ | $L_{\max}$ |
| --- | --- | --- | --- | --- | --- |
| A – Extremely cold | 0.65 | 0.73 | 0.00 | 28.00 | 0.008 |
| B – Hot | $\approx 0.00$ | $\approx 0.00$ | 42.00 | 28.00 | 0.042 |
| C – Cold | 0.64 | 0.73 | 0.11 | 28.00 | 0.020 |
| D – Cold (high $a_r$) | 0.65 | 0.75 | 0.09 | 28.00 | 0.020 |

**Table 3:** tumor-burden metrics under the alternative parameterization. Columns (a)–(d) correspond to the four TIME scenarios: (a) extremely cold; (b) hot; (c) cold; (d) cold with high resistant growth. *X*_0_ denotes the total tumor load at the start of therapy (*t* = 399 days), *X*_min_ the minimum value during the treatment window (399–609 days), and *X*(*t_f_*) the final load at *t_f_* = 700 days. Percentage changes are relative to *X*_0_.

| Metric | (a) | (b) | (c) | (d) |
| --- | --- | --- | --- | --- |
| $X_0$ | 1.07 | 0.96 | 1.07 | 1.07 |
| $X_{\min}$ | 0.65 | 0.00 | 0.64 | 0.65 |
| $X(t_f)$ | 0.73 | 0.00 | 0.72 | 0.75 |
| $\Delta X_{\min}$ [%] | −39.7 | −100.0 | −39.9 | −39.3 |
| $\Delta X_{t_f}$ [%] | −31.7 | −100.0 | −32.3 | −30.3 |

**Table 4:** Effector-lymphocyte and dosing metrics under the alternative parameterization. Columns (a)–(d) correspond to the four TIME scenarios: (a) extremely cold; (b) hot; (c) cold; (d) cold with high resistant growth. *L*_0_ is the effector level at the start of therapy, *L*_min_ the on-treatment nadir, and *L*(*t_f_*) the final value. Percentage changes are relative to *L*_0_. Total ICI and chemotherapy doses are reported as integrated exposure over the 10 treatment intervals (units consistent with the model).

| Metric | (a) | (b) | (c) | (d) |
| --- | --- | --- | --- | --- |
| $L_0$ | 0.004 | 0.035 | 0.015 | 0.015 |
| $L_{\min}$ | 0.003 | 0.023 | 0.015 | 0.015 |
| $L(t_f)$ | 0.002 | 0.012 | 0.020 | 0.020 |
| $\Delta L_{\min}$ [%] | −32.6 | −34.9 | −2.2 | −2.6 |
| $\Delta L_{t_f}$ [%] | −42.2 | −66.0 | +34.2 | +29.7 |
| ICI dose | 0.00 | 42.00 | 0.11 | 0.09 |
| Chemo dose | 28.00 | 28.00 | 28.00 | 28.00 |

#### (i) Extremely cold TIME (case a)

In the extremely cold TIME, the optimizer again con-verges to a chemotherapy-only strategy with *v_I_*(*t*) ≡ 0 throughout the treatment horizon (Fig. 7a). Chemotherapy is delivered at relatively high rates during the first two intervals and then rapidly tapered to zero, in order to respect the cumulative dose cap of 28 units (Table. 4). This schedule yields a transient debulking from *X*_0_ = 1.07 at *t* = 399 days to a nadir of *X*_min_ = 0.65, corresponding to a 39.7% reduction (Table. 3). After treatment cessation, the tumor slowly regrows and reaches *X*(*t_f_*) = 0.73 at day 700, a net 31.7% reduction relative to the start of therapy

The effector compartment remains severely compromised. Effector lymphocytes decrease monotonically from *L*_0_ = 0.004 to *L*(*t_f_*) = 0.002, with an on-treatment nadir of *L*_min_ = 0.003, representing a 32.6% drop during therapy and an overall 42.2% loss by the end of follow-up (Table. 4). Thus, in an extremely cold TIME, the optimizer continues to abandon ICI, achieving only modest tumor shrinkage while further eroding an already fragile immune compartment.

#### (ii) Hot TIME (case b)

Under the alternative parameterization, the hot TIME still supports aggressive combination therapy. The optimizer again saturates both the ICI (*S_I_* = 42) and chemotherapy (*S_M_* = 28) dose budgets, but the scheduling of pulses is slightly adjusted compared with case 1 to account for the increased penalty on drug burden. Tumor burden falls from *X*_0_ = 0.96 to *X*_min_ = 0 during the treatment window and remains at *X*(*t_f_*) = 0 at day 700, corresponding to complete and durable remission (Table. 3).

Effector lymphocytes initially expand but then contract substantially, reflecting the combined impact of chemotherapy-induced lymphodepletion and sustained antigen clearance. *L*(*t*) decreases from *L*_0_ = 0.035 to *L*(*t_f_*) = 0.012, a 66.0% reduction (Table. 4). In this phenotype, the framework willingly “spends” both cytotoxic and immune resources to secure cure, highlighting a clinically familiar trade-off between curative intent and long-term immune reserve.

#### (iii) Cold TIME (case c)

In the moderately cold TIME, the optimizer identifies a compromise schedule. Chemotherapy is again used to its full extent (*S_M_* = 28), but ICI exposure is reduced to a negligible level (*S_I_* = 0.11, three orders of magnitude smaller than in the hot case), deployed as a brief, low-intensity adjunct rather than a primary driver of control (Fig. 7c). Tumor burden decreases from *X*_0_ = 1.07 to *X*_min_ = 0.64 during therapy (39.9% reduction) and stabilizes at *X*(*t_f_*) = 0.72 at day 700 (32.3% reduction; Table. 3).

Unlike the extremely cold phenotype, however, effector lymphocytes are preserved or slightly improved. *L*_min_ remains essentially unchanged relative to baseline (Δ*L*_min_ = −2.2%), and *L*(*t_f_*) = 0.020 exceeds *L*_0_ by 34.2% (Table. 4). This scenario illustrates that, when some immune competence is present but ICI responsiveness is limited, the optimizer gravitates toward chemotherapy-dominated regimens that protect and modestly expand the effector pool.

#### (iv) Cold TIME with high antigen release (case d)

The cold TIME with elevated resistant-cell proliferation *a_r_* exhibits similar qualitative behavior but slightly worse tumor control than case c. The chemotherapy schedule and total dose remain practically unchanged (*S_M_* = 28), whereas ICI usage is further reduced to a total of 0.09 units (Table. 4). Tumor burden falls from *X*_0_ = 1.07 to *X*_min_ = 0.65 during the treatment window (39.3% reduction) and rebounds to *X*(*t_f_*) = 0.75 by day 700, corresponding to a net 30.3% reduction (Table. 3), slightly inferior to case c.

Despite the heightened malignant growth, the effector dynamics remain favorable: the nadir *L*_min_ deviates only marginally from baseline (Δ*L*_min_ = −2.6%), and *L*(*t_f_*) = 0.020 ends 29.7% above *L*_0_ (Table. 4). Here, the optimizer again chooses to preserve immune capacity at the expense of accepting a somewhat higher residual tumor burden.

#### (v) Summary of parameter effects

Across the four TIMEs, the alternative preference weighting reveals a clear pattern. In the hot, highly ICI-responsive microenvironment, the optimizer continues to exhaust both dose budgets, achieving complete tumor eradication at the cost of substantial effector contraction. In the extremely cold TIME, even after reweighting the objective, ICI remains unused and chemotherapy alone delivers only partial, transient control coupled with further immune suppression. In the intermediate cold scenarios, the framework discovers compromise regimens: chemotherapy is fully deployed, ICI is used at vanishingly small doses, and the resulting schedules deliver moderate debulking while preserving or even augmenting effector lymphocytes. Taken together with the identical-parameter experiments, these results show that the proposed dynamic optimization framework reallocates limited chemo–ICI resources across TIME phenotypes in a clinically interpretable way, exposing explicit trade-offs between tumor debulking and long-term immune preservation.

## 5 Conclusion

We presented a continuous-time optimal control framework that couples a mechanistic tumor–immune model with constrained dynamic optimization to explore how microenvironmental context shapes chemotherapy–ICI combination strategies. By explicitly representing sensitive and resistant tumor subpopulations, effector CD8^+^ T cells and drug concentrations within a six-dimensional ODE system, and by enforcing cumulative dose, path and ramp constraints, the framework yields dosing schedules that are both numerically tractable and clinically interpretable. Across four representative TIME phenotypes, the optimizer autonomously adapts the timing and intensity of therapy, suppressing futile ICI use in extremely cold TIMEs, exploiting strong chemo–immunotherapy synergy in hot TIMEs, and identifying intermediate, chemotherapy-dominated regimens in cold TIMEs with limited or partially preserved immune competence.

These optimized regimens quantify trade-offs between tumor control, drug burden and effector preservation and complement more flexible but less transparent reinforcement learning approaches. At the same time, the current model deliberately simplifies toxicity, pharmacokinetics, spatial heterogeneity and patient-to-patient variability, and it is only calibrated to a small set of illustrative TIME parameterizations. Extending the framework to richer pharmacological descriptions, virtual patient populations and calibration against longitudinal clinical or high-dimensional TIME data will be essential for quantitative prediction. Nonetheless, the proposed dynamic optimization module provides a reusable computational building block for deriving phenotype-specific, constraint-aware dosing strategies from tumor–immune models and a starting point for integrating mechanistic optimization into in silico trial design and quantitative systems pharmacology pipelines.

## 6 Competing interests

The authors declare no competing interests.

## 7 Author contributions statement

Kai Gong conducted the modeling, algorithm development, numerical experiments. Tong Lu reviewed the manuscript from his clinical perspective. Xu Wang checked the data and figures. Xing-gao Liu supervised the project.

## Notes

### Competing Interest Statement

The authors have declared no competing interest.

### Funding

- National Natural Science Foundation of China: 62073288, 12075212

