## Supplement Text for "Dynamic optimization of chemo–immunotherapy sequencing reveals phenotype-dependent regimens across tumor immune microenvironments"

#### 1 Restatement of the tumor-immune-therapy model

For completeness, we summarize the ordinary differential equation (ODE) system used in the dynamic optimization framework. The six-dimensional augmented state is

$$x(t) = \big(X_s(t), X_r(t), L(t), I(t), M(t), J(t)\big)^\top \tag{S1}$$

where $X_s(t)$ denotes the burden of treatment-sensitive tumor cells, $X_r(t)$ denotes the burden of resistant tumor cells, $L(t)$ denotes effector CD8$^+$ T cells in the tumor or peripheral compartment, $I(t)$ denotes the concentration of ICI, and $M(t)$ denotes the concentration of chemotherapy, and $J$ accumulates the running cost. The control vector is $u(t) = (v_I(t), v_M(t))$, representing the effective dosing rates of ICI and chemotherapy. The total tumor burden is $X(t) = X_s(t) + X_r(t)$.

The tumor-immune-therapy dynamics are

$$\dot{X}_s = a_s X_s \left(1 - \frac{X_s + \alpha_{sr} X_r}{X_{\max}}\right) - D_L X_s - D_M X_s \tag{S2}$$

$$\dot{X}_r = a_r X_r \left(1 - \frac{X_r + \alpha_{rs} X_s}{X_{\max}}\right) - D_L X_r \tag{S3}$$

$$\dot{L} = -m_L L + r\Phi(X, D_L, D_M) L \left(1 - \frac{L}{L_{\max}}\right) - h_L \eta_{RL} L - q\frac{X}{X + u_{\text{bio}}} L - K_L(1 - e^{-M}) L \tag{S4}$$

$$\dot{I} = -\gamma_I I + v_I \tag{S5}$$

$$\dot{M} = -\gamma_M M + v_M \tag{S6}$$

$$\dot{J} = w_2(X_s + X_r) + w_3 v_M + w_4 v_I - w_1 L \tag{S7}$$

The intermediate quantities encoding checkpoint inhibition and chemo-immune interactions are given by

$$\mu = \mu_0(1 - I)(1 + p(1 - e^{-M})) \tag{S8}$$

$$Z = \frac{L(1-\mu)}{X} \tag{S9}$$

$$D_L = d\,\frac{Z^\ell}{s + Z^\ell} \tag{S10}$$

$$D_M = K_{Xs}\,(1 - e^{-M}) \tag{S11}$$

$$\tag{S12}$$

where $d > 0$ and $\ell > 0$ control the shape of immune-mediated killing. $\mu_0$ is the baseline checkpoint suppression strength, and $K_{Xs}$ scales chemo-induced killing. Effector stimulation is modeled through a composite signal

$$\Phi(X, D_L, D_M) = \frac{j_C X}{k_C + X} + \frac{j_L D_L X}{k_L + D_L X} + \frac{j_M D_M X_s}{k_M + D_M X_s} \tag{S13}$$

where $X = X_s + X_r$ and the three terms represent baseline antigenic stimulation, CD8$^+$ enhanced immunogenic cell death, and chemo-induced immunogenicity in the sensitive tumor compartment.

### 2 Representative analytical derivatives for the Jacobian $A(x, u) = \partial F/\partial x$

The Jacobian $A(x, u)$ of the right-hand side $F(x, u; \theta)$ with respect to the state variables is required for the embedded sensitivity equations. Here we list representative entries; the remaining entries follow by analogous application of the chain rule.

#### 2.1 Useful partial derivatives

We define the logistic saturation terms

$$\Phi_s = \frac{X_s + \alpha_{sr} X_r}{X_{\max}}, \;\; \Phi_r = \frac{X_r + \alpha_{rs} X_s}{X_{\max}} \tag{S14}$$

Their derivatives are

$$\frac{\partial \Phi_s}{\partial X_s} = \frac{1}{X_{\max}}, \;\; \frac{\partial \Phi_s}{\partial X_r} = \frac{\alpha_{sr}}{X_{\max}}, \;\; \frac{\partial \Phi_r}{\partial X_r} = \frac{1}{X_{\max}}, \;\; \frac{\partial \Phi_r}{\partial X_s} = \frac{\alpha_{rs}}{X_{\max}} \tag{S15}$$

The quantity $Z = L(1 - \mu)/X$ satisfies

$$\frac{\partial Z}{\partial X_s} = -\frac{L(1-\mu)}{X^2}, \quad \frac{\partial Z}{\partial X_r} = -\frac{L(1-\mu)}{X^2} \tag{S16}$$

$$\frac{\partial Z}{\partial L} = \frac{1-\mu}{X}, \quad \frac{\partial Z}{\partial I} = -\frac{L}{X}\frac{\partial \mu}{\partial I}, \quad \frac{\partial Z}{\partial M} = -\frac{L}{X}\frac{\partial \mu}{\partial M} \tag{S17}$$

with

$$\frac{\partial \mu}{\partial I} = -\mu_0\Big[1 + p(1 - e^{-M})\Big], \quad \frac{\partial \mu}{\partial M} = \mu_0(1 - I)pe^{-M} \tag{S18}$$

The derivatives of $D_L$ and $D_M$ with respect to a generic variable $y \in \{X_s, X_r, L, I, M\}$ are

$$\frac{\partial D_L}{\partial y} = d\,\ell Z^{\ell-1}\frac{\partial Z}{\partial y} + \ell Z^{\ell-1}\frac{\partial Z}{\partial y} \tag{S19}$$

$$\frac{\partial D_M}{\partial y} = K_{Xs}e^{-M}\frac{\partial M}{\partial y} = \begin{cases} 0, & y \in \{X_s, X_r, L, I\}, \\ -K_{Xs}e^{-M}, & y = M. \end{cases} \tag{S20}$$

Derivatives of $U$ with respect to $X_s, X_r, D_L, D_M$ can be obtained by straightforward differentiation of the rational terms; we omit the full expressions here as they are long but mechanically derived.

#### 2.2 Tumor equations

The sensitive tumor equation is

$$f_{X_s} = a_s X_s(1 - \Phi_s) - X_s(D_L + D_M) \tag{S21}$$

Differentiating with respect to the state variables yields

$$\frac{\partial f_{X_s}}{\partial X_s} = a_s(1-\Phi_s) - a_s X_s \frac{\partial \Phi_s}{\partial X_s} - (D_L + D_M) - X_s \left(\frac{\partial D_L}{\partial X_s} + \frac{\partial D_M}{\partial X_s}\right) \tag{S22}$$

$$\frac{\partial f_{X_s}}{\partial X_r} = -a_s X_s \frac{\partial \Phi_s}{\partial X_r} - X_s \left(\frac{\partial D_L}{\partial X_r} + \frac{\partial D_M}{\partial X_r}\right) \tag{S23}$$

$$\frac{\partial f_{X_s}}{\partial L} = -X_s \left(\frac{\partial D_L}{\partial L} + \frac{\partial D_M}{\partial L}\right) \tag{S24}$$

$$\frac{\partial f_{X_s}}{\partial I} = -X_s \left(\frac{\partial D_L}{\partial I} + \frac{\partial D_M}{\partial I}\right) \tag{S25}$$

$$\frac{\partial f_{X_s}}{\partial M} = -X_s \left(\frac{\partial D_L}{\partial M} + \frac{\partial D_M}{\partial M}\right) \tag{S26}$$

The resistant tumor equation is

$$f_{X_r} = a_r X_r (1-\Phi_r) - D_L X_r \tag{S27}$$

Derivatives are

$$\frac{\partial f_{X_r}}{\partial X_r} = a_r(1-\Phi_r) - a_r X_r \frac{\partial \Phi_r}{\partial X_r} - D_L - X_r \frac{\partial D_L}{\partial X_r} \tag{S28}$$

$$\frac{\partial f_{X_r}}{\partial X_s} = -a_r X_r \frac{\partial \Phi_r}{\partial X_s} - X_r \frac{\partial D_L}{\partial X_s} \tag{S29}$$

$$\frac{\partial f_{X_r}}{\partial L} = -X_r \frac{\partial D_L}{\partial L} \tag{S30}$$

$$\frac{\partial f_{X_r}}{\partial I} = -X_r \frac{\partial D_L}{\partial I} \tag{S31}$$

$$\frac{\partial f_{X_r}}{\partial M} = -X_r \frac{\partial D_L}{\partial M} \tag{S32}$$

#### 2.3 Immune, drug, and cost equations

The immune equation can be written as

$$f_L = -m_L L + r\Phi(X, D_L, D_M) L \left(1 - \frac{L}{L_{\max}}\right) - h_L \eta_{RL} L - q\frac{X}{X + u_{\text{bio}}} L - K_L(1 - e^{-M}) L \tag{S33}$$

Its derivatives involve products and compositions of $U$, $D_L$, and $D_M$. For example,

$$\frac{\partial f_L}{\partial L} = -m_L + r\Phi \left(1 - \frac{2L}{L_{\max}}\right) + rL\left(1 - \frac{L}{L_{\max}}\right)\frac{\partial \Phi}{\partial L} - h_L \eta_{RL} - \frac{qX}{X + u_{\text{bio}}} - K_L(1 - e^{-M}) \tag{S34}$$

and

$$\frac{\partial f_L}{\partial X_s} = r\,L\left(1 - \frac{L}{L_{\max}}\right)\frac{\partial \Phi}{\partial X_s} - q\frac{\partial}{\partial X_s}\left(\frac{X}{X + u_{\text{bio}}}\right) L, \tag{S.35}$$

with $\partial f_L / \partial X_r$ obtained analogously. Terms involving $\partial \Phi / \partial D_L$ and $\partial \Phi / \partial D_M$ can be further expanded using S19–S20.

The drug and cost equations have simple linear structure:

$$f_I = -\gamma_I I + v_I \tag{S35}$$

$$f_M = -\gamma_M M + v_M \tag{S36}$$

$$f_J = w_2(X_s + X_r) + w_3 v_M + w_4 v_I - w_1 L \tag{S37}$$

Hence

$$\frac{\partial f_I}{\partial I} = -\gamma_I, \ \frac{\partial f_I}{\partial v_I} = 1, \ \frac{\partial f_M}{\partial M} = -\gamma_M, \ \frac{\partial f_M}{\partial v_M} = 1 \tag{S38}$$

$$\frac{\partial f_J}{\partial X_s} = w_2, \ \frac{\partial f_J}{\partial X_r} = w_2, \ \frac{\partial f_J}{\partial L} = -w_1, \ \frac{\partial f_J}{\partial v_M} = w_3, \ \frac{\partial f_J}{\partial v_I} = w_4 \tag{S39}$$

#### 2.4 Embedded sensitivity equations and parameter injection

Let $P = 2N$ denote the number of control parameters. We collect these in a parameter vector

$$p = (v_{I,0}, v_{M,0}, \dots, v_{I,N-1}, v_{M,N-1})^\top \in \mathbb{R}^P \tag{S40}$$

For each time $t$ we define the sensitivity matrix

$$S(t) = \frac{\partial x(t)}{\partial p} \in \mathbb{R}^{6\times P} \tag{S41}$$

The sensitivity dynamics follow the standard linearized form

$$\dot{S}(t) = A(x(t), u(t))S(t) + B(x(t), u(t))\frac{\partial u(t)}{\partial p} \tag{S42}$$

where $A = \partial F/\partial x$ is the Jacobian described in Section S2 and $B = \partial F/\partial u$ collects the partial derivatives of $F$ with respect to $(v_I, v_M)$.

Within the $k$-th treatment interval $[t_k, t_{k+1})$, the control is constant:

$$u(t) = u_k = (v_{I,k}, v_{M,k})^\top \tag{S43}$$

We realize the mapping $\partial u/\partial p$ via a sparse injection matrix $E_k \in \mathbb{R}^{2\times P}$ that selects the two relevant parameters:

$$\frac{\partial u(t)}{\partial p} = \begin{cases} E_k, & t \in [t_k, t_{k+1}), \\ 0, & \text{otherwise}, \end{cases} \tag{S44}$$

where $E_k$ has ones in the columns corresponding to $v_{I,k}$ and $v_{M,k}$ and zeros elsewhere. In the implementation, this is equivalent to using a one-hot vector $\delta \in \mathbb{R}^P$ for each subsegment.

For numerical integration, we concatenate the state and sensitivities into a single augmented state

$$y(t) = \big(x(t), \mathrm{vec}(S(t))\big)^\top \in \mathbb{R}^{6+6P} \tag{S45}$$

and integrate the ODE system

$$\frac{d}{dt}\begin{bmatrix} x(t) \\ \mathrm{vec}(S(t)) \end{bmatrix} = \begin{bmatrix} F(x(t), u(t); \theta) \\ \mathrm{vec}\Big(A(x(t), u(t))S(t) + B(x(t), u(t))\frac{\partial u}{\partial p}\Big) \end{bmatrix} \tag{S46}$$

##### 2.5 Objective gradient and permutation to the solver's decision vector

Because the running cost has been absorbed into the auxiliary state $J(t)$ via S7, the performance index for a given parameter vector $p$ is

$$J(p) = J(t_f; p) = J(t_f) \tag{S47}$$

The gradient of $J$ with respect to $p$ is directly obtained from the sensitivities at final time:

$$\frac{\partial J}{\partial p} = e_J^\top S(t_f) \tag{S48}$$

where $e_J = (0, 0, 0, 0, 0, 1)^\top$ is the unit vector selecting the sixth component of the state. In practice, this corresponds to taking the last row of $S(t_f)$.

##### 2.6 Jacobians of inequality constraints

We recall the three families of inequality constraints in the dynamic optimization framework.

- cumulative dose caps on ICI and chemotherapy,
- pharmacokinetic safety caps on $I$ and $M$ at the end of each treatment interval,
- ramp constraints on changes in per-interval doses.

###### 2.6.1 Cumulative dose constraints

The cumulative ICI and chemotherapy doses are

$$S_I(p) = \sum_{k \in K_I} v_{I,k}\, \Delta t, \qquad S_M(p) = \sum_{k \in K_M} v_{M,k}\, \Delta t \tag{S49}$$

and the corresponding constraints are

$$g_{\text{dose}}(p) = \begin{pmatrix} D_I^{\text{cap}} - S_I(p) \\ D_M^{\text{cap}} - S_M(p) \end{pmatrix} \geq 0. \tag{S50}$$

The Jacobian of $g_{\text{dose}}$ with respect to the dosing vector $p$ is constant and given by

$$\frac{\partial g_{\text{dose}}}{\partial p} = -\begin{pmatrix} \Delta t\, \chi_{K_I}(0) & 0 & \cdots & \Delta t\, \chi_{K_I}(N-1) & 0 \\ 0 & \Delta t\, \chi_{K_M}(0) & \cdots & 0 & \Delta t\, \chi_{K_M}(N-1) \end{pmatrix} \tag{S51}$$

where $\chi_{K_I}(k)$ and $\chi_{K_M}(k)$ are indicator functions that equal one if the index $k$ belongs to the ICI or chemotherapy treatment window, respectively, and zero otherwise.

###### 2.6.2 Pharmacokinetic safety constraints

Let $I_k$ and $M_k$ denote the values of $I$ and $M$ at time $t_{k+1}$ obtained from the forward integration under a given control vector $p$, and let $S_I(t_{k+1}, :)$ and $S_M(t_{k+1}, :)$ denote the corresponding rows of the sensitivity matrix

$$S(t) = \frac{\partial x(t)}{\partial p}. \tag{S52}$$

The pharmacokinetic safety constraints are

$$g_{\text{path}}(p) = \left(I^{\max} - I_k,\ M^{\max} - M_k\right)_{k=0}^{N-1} \geq 0. \tag{S53}$$

Their Jacobian rows are obtained directly from the sensitivities:

$$\frac{\partial}{\partial p}(I^{\max} - I_k) = -S_I(t_{k+1}, :), \qquad \frac{\partial}{\partial p}(M^{\max} - M_k) = -S_M(t_{k+1}, :) \tag{S54}$$

where $S_I$ and $S_M$ denote the rows of $S$ corresponding to the $I$ and $M$ components of the state.

##### 2.6.3 Ramp constraints

For $k = 1, \ldots, N-1$, the ramp constraints on ICI and chemotherapy doses are

$$|v_{I,k} - v_{I,k-1}| \le \rho_I, \qquad |v_{M,k} - v_{M,k-1}| \le \rho_M \tag{S55}$$

We implement them as two-sided linear inequalities,

$$\begin{aligned} g^{+}_{I,k}(p) &= \rho_I - (v_{I,k} - v_{I,k-1}) \ge 0, \\ g^{-}_{I,k}(p) &= \rho_I + (v_{I,k} - v_{I,k-1}) \ge 0 \end{aligned} \tag{S56}$$

and similarly for $g^{\pm}_{M,k}(p)$. The nonzero entries of their Jacobians are

$$\frac{\partial g^{+}_{I,k}}{\partial v_{I,k}} = -1, \quad \frac{\partial g^{+}_{I,k}}{\partial v_{I,k-1}} = 1, \qquad \frac{\partial g^{-}_{I,k}}{\partial v_{I,k}} = 1, \quad \frac{\partial g^{-}_{I,k}}{\partial v_{I,k-1}} = -1 \tag{S57}$$

and analogously for the chemotherapy variables with $\rho_M$. All other entries are zero, so the ramp-constraint Jacobian is sparse and time-independent.
